# Artifact-Minimized High-Ratio Image Compression with Preserved Analysis Fidelity

**DOI:** 10.1101/2024.07.17.603794

**Authors:** Bin Duan, Logan A Walker, Bin Xie, Wei Jie Lee, Alexander Lin, Yan Yan, Dawen Cai

## Abstract

Recent advances in microscopy have pushed imaging data generation to an unprecedented scale. While scientists benefit from higher spatiotemporal resolutions and larger imaging volumes, the increasing data size presents significant storage, visualization, sharing, and analysis challenges. Lossless compression typically reduces the data size by <4 fold, whereas lossy compression trades smaller data size for the loss of a precise reconstruction of the original data. Here, we develop a novel quantization method and an artifact metric for automated compression parameter optimization that preserves information fidelity. We show that, when combined with the AV1 video codec, we achieve tens to ten thousand folds of data compression while introducing negligible visual defects or quantification errors in single-molecule localization and segmentation analyses. We developed an HDF5 filter with FFMPEG library support for convenient community adaptation. For instance, HDF5-enabled ImageJ plugins can now be seamlessly extended to support AV1 compression and visualization to handle terabyte-scale images.

---

Advances in modern imaging modalities have pushed data generation into the terabyte-to-petabyte scale^1–3^. This results in tremendous costs for data storage and distribution, especially when frequently transferring data via the Internet for remote collaboration. Lossless compression has been implemented to reduce the data size while preserving all information^4–6^, however, it typically only achieves, at most, a 4-fold reduction^6^ for biomedical images. Lossy compression, on the other hand, selectively discards information to achieve higher compression ratios at the expense of analysis fidelity^3,7^ due to artifacts generated in its quantization and compression steps. Despite a loss of image quality, recent work has shown the potential of using lossy compression for quantitative microscopy experiments^3,7–10^. For example, to meet the real-time data transfer needs for a high-throughput super-resolution localization microscope, a lossy data quantization step was first applied to the raw images to reduce the dynamic range from 16-bit to *<*8-bit^11^. The quantized data stream was further losslessly encoded with a Huffman coding scheme^12^ to achieve ∼ 7-fold compression with a 4*%* error rate of recalling single fluorescent spots^3^.

2D biomedical image compression typically applies well-established lossy compression algorithms, such as JPEG^13^, JPEG2000^14^, and, more recently, machine learning^15^. For 3D volumetric image compression, video codecs such as H.264^16^, H.265^17^, and AV1^18^, can be adapted by treating one image axis, typically the *z* axis, as the pseudo-time axis. For example, H.265 has been used to compress 3D fluorescent image data for lightweight and distributed human reconstruction of neuron morphologies from the whole mouse brain^7^. In another example, combining machine learning-based denoising and AV1 compression, a 3D volumetric electron microscopy dataset was reduced by approximately 17 folds while achieving ∼ 99*%* edge accuracy between results obtained from compressed and uncompressed images, although only achieving <90*%* accuracy of synapse detection^9^. These examples reveal that state-of-the-art video compression has great potential to replace lossless image compression in specific quantitative analyses. That said, there is still much more to be understood about applying lossy compression. For instance, better quantization is needed for consequent high-fidelity compression. Video codec settings also strongly influence the compression ratio and quality trade-offs yet are hard to predict and quantify. Finding a solution to automatically determine the optimal settings is essential for user adaptation because it will eliminate the lengthy and subjective trial-and-error process while achieving superior data reduction and analysis fidelity.

Here, we show the development of a novel quantization method to optimally preserve critical information for faithful quantitative analysis. We compare several mainstream video compression algorithms and determine that AV1 is the best algorithm to be coupled with optimal quantization for high-ratio and high-fidelity compression. We introduce a novel metric for evaluating the artifacts generated in lossy image compression and develop an algorithm to determine the optimal compression parameters for AV1. With these developments, AV1 impressively compresses fluorescence and electron microscopy images by tens or even tens of thousands of folds with negligible errors in single-molecule localization or machine learning segmentation analyses, respectively. Finally, to promote community adoption, we created an HDF5 filter with FFMPEG video codec support and an ImageJ/Fiji^19^ plugin for visualization and manipulation of images with automated optimal lossy compression.

## Results

### Non-linear Quantization maximally preserves information while reducing image bit-depth

Modern microscope cameras can record 16-bit dynamic range to better preserve details from dim to bright regions in the image. Before being compressed or computed upon, high bit-depth images are often transformed into low bit-depth images in a quantization step, which introduces rounding errors and, therefore, is a lossy process. This quantization can be compulsory, especially when most advanced video codec implementations only support up to 10-bit compression, with limited support for higher bits^20–22^. The most common quantization is a direct linear transform from 16 to 8 bits, for example, as implemented in ImageJ/Fiji^19^:

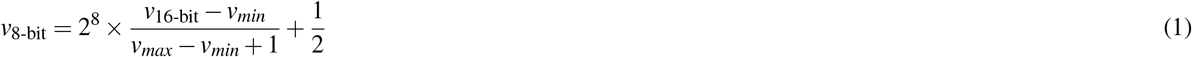

where *v*_*min*_ and *v*_*max*_ represent the minimum and maximum value in an image, respectively. Another popular quantization method is taking the square root for each pixel 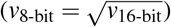, conveniently converting its value from a 2^16^ range into a 2^8^ range. This non-linear quantization method has been recently used to reduce the data transfer bandwidth requirement for space telescope images^11^ and high-throughput localization microscope images^3^.

We have found that linear and square root quantization resulted in losing valuable information in many biomedical images, which were taken under low exposure conditions to minimize sample photobleaching. For instance, an image with a maximum intensity value of 10,000 is considered bright yet still far below the maximum value of 65,535 of a 16-bit detector. Direct application of linear or square root quantization results in the suppression of dynamic range, leading to a higher information loss, particularly in the low-intensity range. Such pixels representing intensity transitions, including noisy backgrounds, require better preservation because they present the most significant contrast to human visual perception and are the focus of many image processing and quantification analyses, including machine learning-based segmentation and computer vision^23,24^. Therefore, we reason that an optimal quantization method should occupy the whole target dynamic range with an emphasis on preserving low-intensity information. In addition, it should be simple and easy to implement to accommodate terabyte-to-petabyte scale image datasets.

To satisfy these criteria, we propose a beta quantization using a generalized power function: *I*_low_ = (*I*_high_)^*β*^, where *I*_low_ and *I*_high_ represent the maximum value of the low-bit and high-bit image, respectively. In this generalized form, canonical square root conversions represent a special case of *β* = 0.5. To compare the fidelity of different quantization methods, we acquired 16-bit confocal images of cell nuclei from a mouse brain section (Fig. 1a). With *I*_high_ = 3, 941, the optimal *β* is calculated as *β* = log 255*/*log 3941 ≈ 0.67, which results in better utilizing the full 8-bit information encoding range after quantization (Fig. 1b-c). To be noted is that high-bit to low-bit quantization always results in an elevation of the low intensity (Fig. 1b), which is equivalent to non-proportionally bias in the encoding bit capacity towards these critical intensity transitions, as mentioned above.

**Fig. 1.**
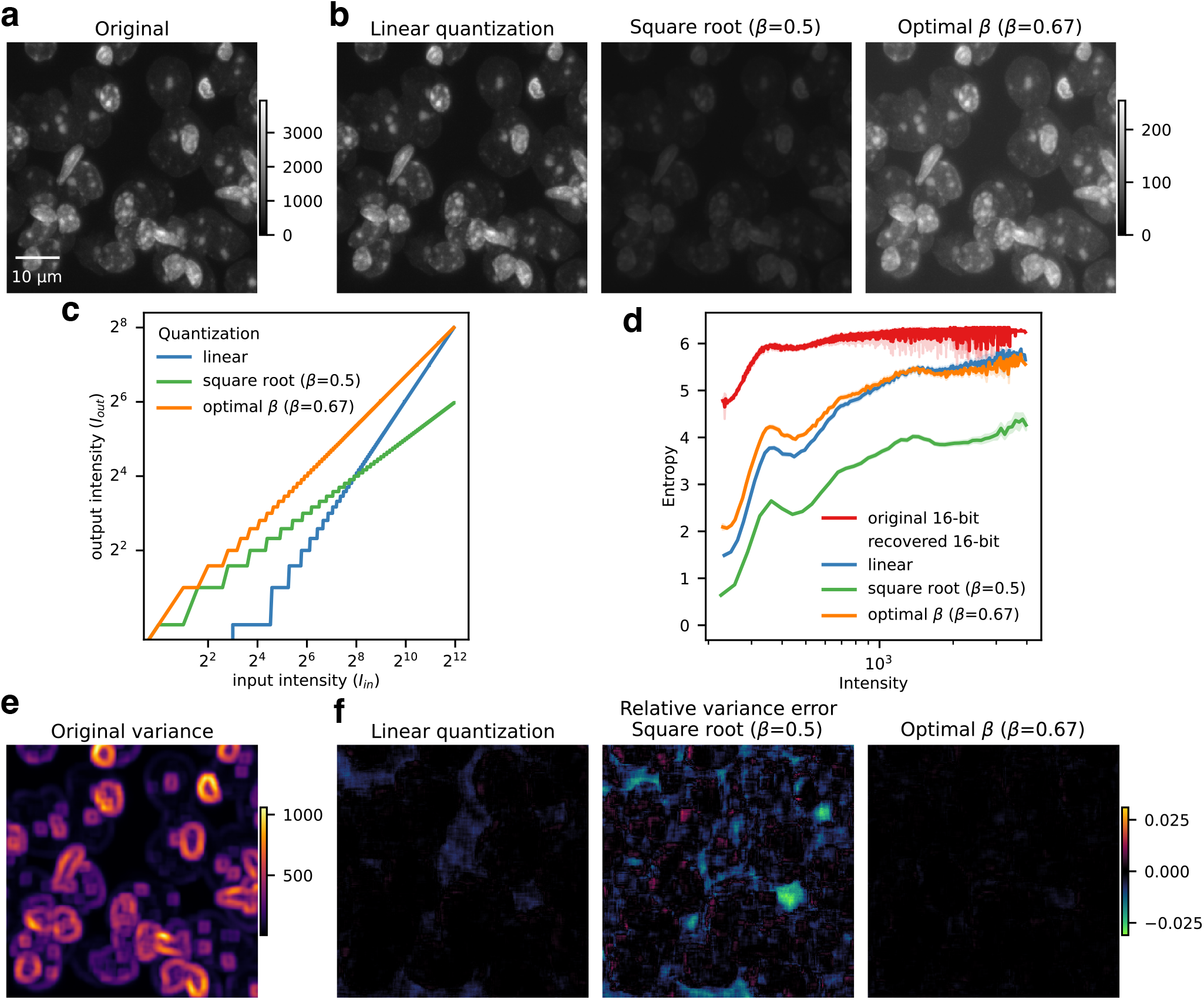
Optimal beta quantization maximally preserves information while reducing image bit-depth. **a**, Original 16-bit confocal images of cell nuclei fluorescently labeled with DRAQ5 from a mouse brain section. **b**, From left to right: 8-bit image transformed by linear quantization, square root quantization (*β* = 0.5), and optimal beta quantization (*β* = 0.67) calculated based on the maximum intensity of the original image (**a**), respectively. **c**, Correspondent input and output intensity plot of different quantization methods. **d**, Intensity-Entropy distribution of the original image **a** and 16-bit images recovered from **b. e**, Intensity variance of the original image **a. f**, From left to right: relative variance errors of the 16-bit images recovered from (**b**), respectively.

Next, we computed the entropy^25^ across the images as an estimation of the information being encoded, where entropy is defined as how many bits need to represent all neighboring values within a 5-pixel radius. Close inspection of the entropy-intensity distribution shows that the raw image has the highest entropies across all intensity values, including the background regions, followed by optimal beta quantization (Fig. 1d, Supplementary Fig. 1). To quantitatively compare different quantization methods in preserving intensity transition details, we computed the difference of relative intensity variance across the whole image. To do so, we first computed the intensity variance in a 5-pixel radius of the original 16-bit image, where a high variance reveals the pixel with rapid intensity change in its surroundings, typically along the boundary of a nucleus (Fig. 1e). Reverse operations were then used to restore the 8-bit quantized images to 16-bit, followed by computing their intensity variance. The relative differences between the intensity variances of each restored image and the original image indicate that optimal beta quantization introduced the least error (Fig. 1f). Taken together, we conclude that our beta quantization method using optimal power transform can better preserve the information encoded in microscopy images, particularly under low light conditions.

### AV1 is a superior lossy video codec for biomedical image compression

Previous studies show that, compared to H.264 and H.265, AV1 has the highest coding efficiency for high-definition gaming and video streaming^26,27^. Here, we perform a benchmarking study that compares pixel-level compression errors, which are essential for reproducible quantitative analyses of biomedical images.

Our benchmark dataset is composed of 5 image samples from different microscopy modalities, including low contrast point scanning confocal microscopy images of NeuN staining^28^, high contrast point scanning confocal images of Brainbow-labeled mouse neurons^29^ (Brainbow), expansion line scanning confocal microscopy images of DRAQ5 stained mouse brain nuclei (DRAQ5), STochastic Optical Reconstruction Microscopy (STORM) images of *β* 2-spectrin labeling in cultured hippocampal neurons^30^, and Electron microscope (EM) scanned Drosophila melanogaster larval neuropil^31^ (Fig. 2, Supplementary Fig. 2). We first benchmarked each video codec’s ability to encode in 8-bit or 10-bit under preset parameter settings that emphasize compression quality by computing the corresponding compression ratio, normalized root mean squared error (NRMSE), peak signal-to-noise ratio (PSNR), and structural similarity index measure (SSIM) (Fig. 2a). We found that they generate high degrees of data preservation with a slight edge going to AV1, while all achieving several fold higher compression ratios compared to the best overall lossless compression method Blosc-Zstd^6,32,33^(Supplementary Fig. 3). In more than ten-fold compression schemes, visual inspection indicates AV1 has the best overall performance, especially when turning on its unique film grain feature under extremely high compression conditions (Fig. 2b-e). However, we also noticed that AV1’s film grain feature introduced extra smoothing, which may be undesirable for single-molecule localization super-resolution imaging applications (Fig. 2d) because it significantly changes the single molecules background noise signatures (detailed below). Nonetheless, we conclude that AV1 is the best among the three most supported video compression codecs due to its overall performance, especially in applications that require and tolerate extremely high compression ratios.

**Fig. 2.**
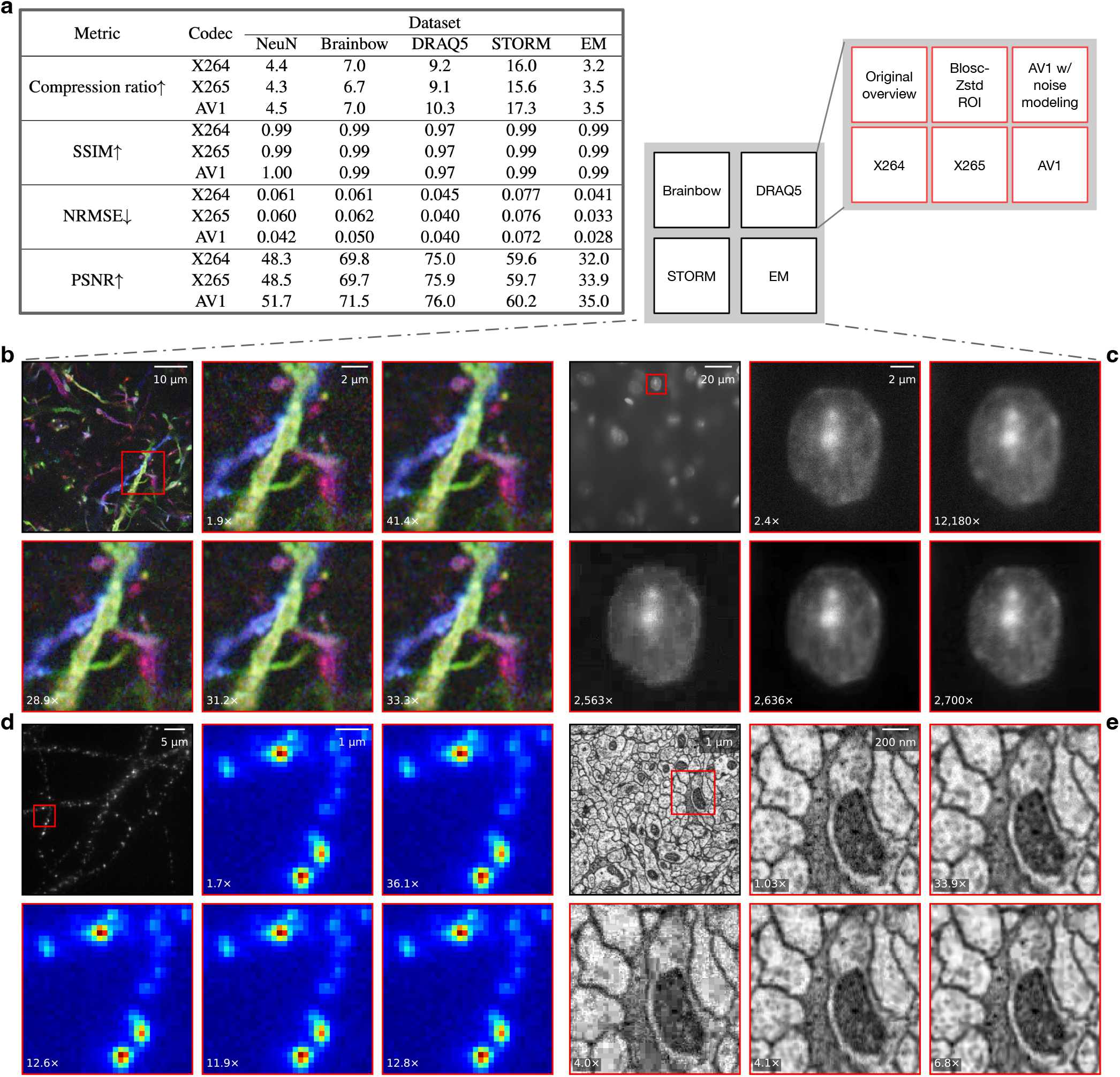
Lossy video compression codec comparison. **a**, Comparison of X264, X265, and AV1 performances using preset parameter settings that emphasize compression quality. The 10-bit or 8-bit compression mode was applied to 16-bit fluorescence or 8-bit EM images, respectively. **b**–**e**, Visual inspection of lossless compression (Blosc-Zstd) and lossy compressions (X264, X265, and AV1) using preset parameter settings that emphasize compression ratio. NRMSE, normalized root mean squared error; PSNR, peak signal-to-noise ratio; SSIM, structural similarity index measure.

### Non-linear quantization with AV1 lossy compression preserves excellent fitting accuracy in single-molecule localization microscopy (SMLM)

SMLM improves roughly ten-fold in spatial resolution over conventional diffraction-limited fluorescence microscopy at the price of acquisition time and data volume^34^. For SMLM techniques, such as STORM, a single field of view with only tens of micrometers in diameter typically requires 10,000 to 100,000 raw camera data frames^35^. Due to the short exposure time, SMLM images are often very noisy, making them notoriously difficult to compress using lossless algorithms. At the same time, local noise signatures should not be significantly altered since estimating background noise crucially affects the accuracy of fitting a point spread function (PSF)^36^. In a recently published high-throughput SMLM platform, a linearly scaled square root transform was first applied to quantize the 16-bit raw images into a bit-depth that is effectively smaller than 8-bit, and then the lossless Zstd compression was applied to reduce the data bandwidth for real-time data transfer needs^3^. As optimal beta transform quantization preserves more information encoded in the low-intensity background-transition pixels (Fig. 1d, 1f), we expect it will lead to higher fitting accuracy in determining the localization of a single molecule PSF. We further hypothesize that using AV1 to compress optimally quantized SMLM data, especially in 10-bit, would introduce negligible fitting errors while achieving higher compression efficiency.

We downloaded a published STORM image dataset^30^ (Fig. 3a), compressed it using various compression strategies and settings, and then decompressed it back to 16-bit images for localization analyses. While the Blosc-Zstd lossless algorithm compresses the raw data by a 1.6× ratio, all lossy compression strategies achieve compression ratios greater than 3.5×. As expected, linear-scaled 8-bit square root quantization followed by Blosc-Zstd lossless compression (sqrt8-LS, Fig. 3b, blue line) resulted in higher compression ratios than 8-bit optimal beta quantization with the same linear scaling factors (beta8-LS, Fig. 3b, orange line). As AV1 supports 10-bit coding, we also used 10-bit optimal beta quantization without linear scaling (beta10) to preserve more background information. Beta10, followed by AV1, generated much higher compression ratios (Fig. 3b, green line). That said, the ultimate evaluation requires comparing how much error each compression introduces when fitting single molecule locations. To do so, we utilized the ThunderSTORM^37^ software to identify fluorescent spots in the decompressed 16-bit images and quantified their centroid localization error (CLE) and spot identification error rate (SIER) corresponding to those identified in the original image (Fig. 3c). As expected, beta8-LS can achieve the least errors (<3.5 nm CLE and <3% SIER) in the lowest, <4× compression range. In the 4-10× compression range, the compression ratio vs. error performance of beta8-LS and sqrt8-LS are extremely similar (<20 nm CLE and <6% SIER). Beta10-AV1 has the best compression vs. error performance if 6-fold data reduction is desired. Interestingly, we found much higher false negative than false positive rates in all compression strategies (Supplementary Fig. 4). That said, we suspect that the missing spots may be retrieved using alternative fitting algorithms^38–40^. Finally, we visually examined the compression quality, focusing on comparing settings that generated the same compression ratios using all three strategies (Fig. 3d-i). We found even at >40× compression, the beta10-AV1 compressed image showed a high degree of similarity to the raw image, which is better than sqrt8-LS at 13× compression (Fig. 3c, 3j).

**Fig. 3.**
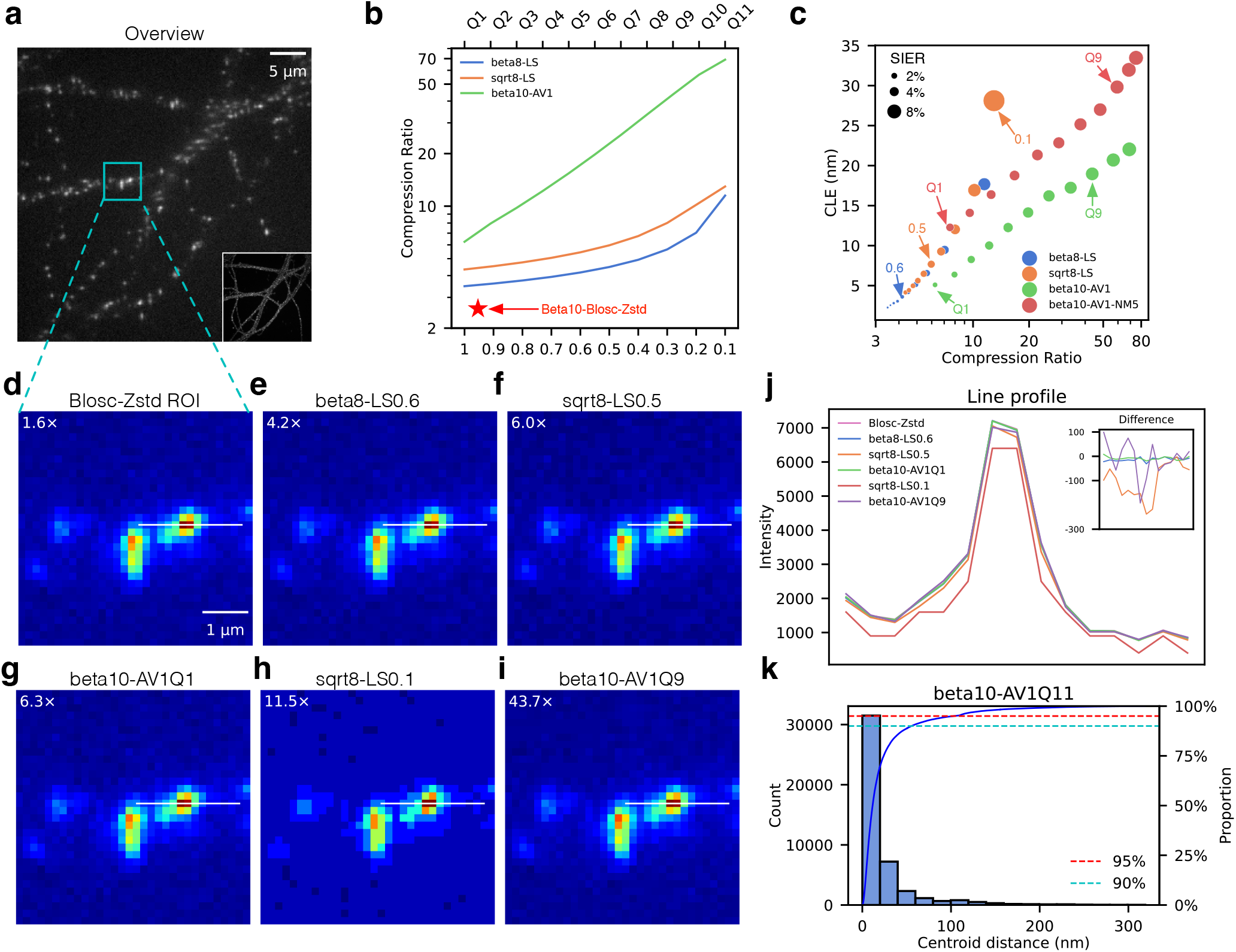
Beta quantization with AV1 lossy compression preserves faithful single-molecule localization analysis. **a**, Single STORM image of *β* 2-spectrin labeling in cultured hippocampal neurons. **b**, Plot of compression ratios achieved using different compression strategies, in which scaling factors of 1 to 0.1 were used in sqrt8-LS and beta8-LS, quality factors of Q1 to Q11 were used in beta10-AV1. **c**, Plot of compression ratio, centroid localization error (CLE), and spot identification error rate (SIER). **d**–**i**, Zoom-in of the cyan box in (**a**) compressed by indicated strategies. **j**, Intensity line profile of white line in (**d**-**i**). Inset shows the pixel intensity difference between the original image (**a**) and images being lossy compressed (**e, f, g, i**) along the white line. sqrt8-LS, linear-scaled 8-bit square root quantization followed by Blosc-Zstd lossless compression; beta8-LS, linear-scaled 8-bit optimal beta quantization followed by Blosc-Zstd lossless compression; beta10-AV1, 10-bit optimal beta quantization followed by AV1 lossy compression. **k**, Histogram and cumulative proportion plots of centroid distances between the fitting spots of the uncompressed image and beta10-AV1Q11 compressed image.

One common application of SMLM is to reconstruct super-resolution images using the fitted centroids of the identified spots from the fluorescent images (Supplementary Fig. 5). To evaluate the impact of image compression on this application, we calculated the degree of agreement, with a 95% confidence interval, between the centroids fitted using raw images and those fitted using compressed images. We found that sqrt8-LS can compress to 5.6× while maintaining a 99% agreement, and an 8.8× data reduction can be achieved if lowering the agreement criteria to 96%, which agrees with recently published results^3^. Beta10-AV1 improves the compression to 14.5× or 69× if a 99% or 96% fitting agreement is required, respectively. SMLM is also commonly used to decode gene barcodes in spatial transcriptomics^41–43^. For example, in the MERFISH^41^ protocol, a 100 nm distance threshold is normally set to allocate fluorescent spots imaged in consecutive rounds as the same mRNA molecule^44^. As shown in Fig. 3c, beta10-AV1Q2 can achieve an 8× data compression while having a 2% false positive/false negative total error rate (TER) and 6 nm mean localization error relative to fittings using raw images. With a higher error tolerance, beta10-AV1Q11 can achieve a 69× compression while having an 8% TER with over 95% fitted spots are below the 100 nm MERFISH error thresholds^44^ (Fig. 3k).

### Automated determination of noise modeling parameters minimizes lossy compression artifacts

All lossy compression codecs display various artifacts, including lower contrast, blurry, and, most prominently, the so-called block artifact^45^ when reaching very high compression ratios (Fig. 4a). The block artifact appears as a mosaic pattern that breaks the image into discontinuous blocks, each of which corresponds to a non-overlapping window that the compression statistics approximation is computed in parallel to reduce computation complexity and increase scalability^16–18^. High-ratio compression causes a severe block artifact because of information loss within each window. As a biomedical image can be decomposed into a noise background and a feature foreground, reducing background noise during acquisition can effectively improve image compression ratios^46^. Coupling post-acquisition denoising with less aggressive settings, video codecs can also achieve more faithful compression for segmentation analysis^9^. Interestingly, SVT-AV1 and other AV1 codecs include a film grain synthesis (FGS) feature, which is fundamentally a denoise-and-noise modeling process followed by a noise synthesis process to preserve natural-looking noise in the decompressed video stream^47^ (Fig. 4b). We noted that AV1 FGS could effectively reduce the block artifact with a trade-off of generating a slightly blurred image, even with a *>*10,000× compression (Fig. 4a). In practice, the compression quality is affected by both the constant rate factor (Q), and the noise modeling factor (NM), two positive integers that determine the desired bitrate and the noise modeling level, respectively. However, determining the proper parameter values is an empirical process, which could be time-consuming and non-objective.

**Fig. 4.**
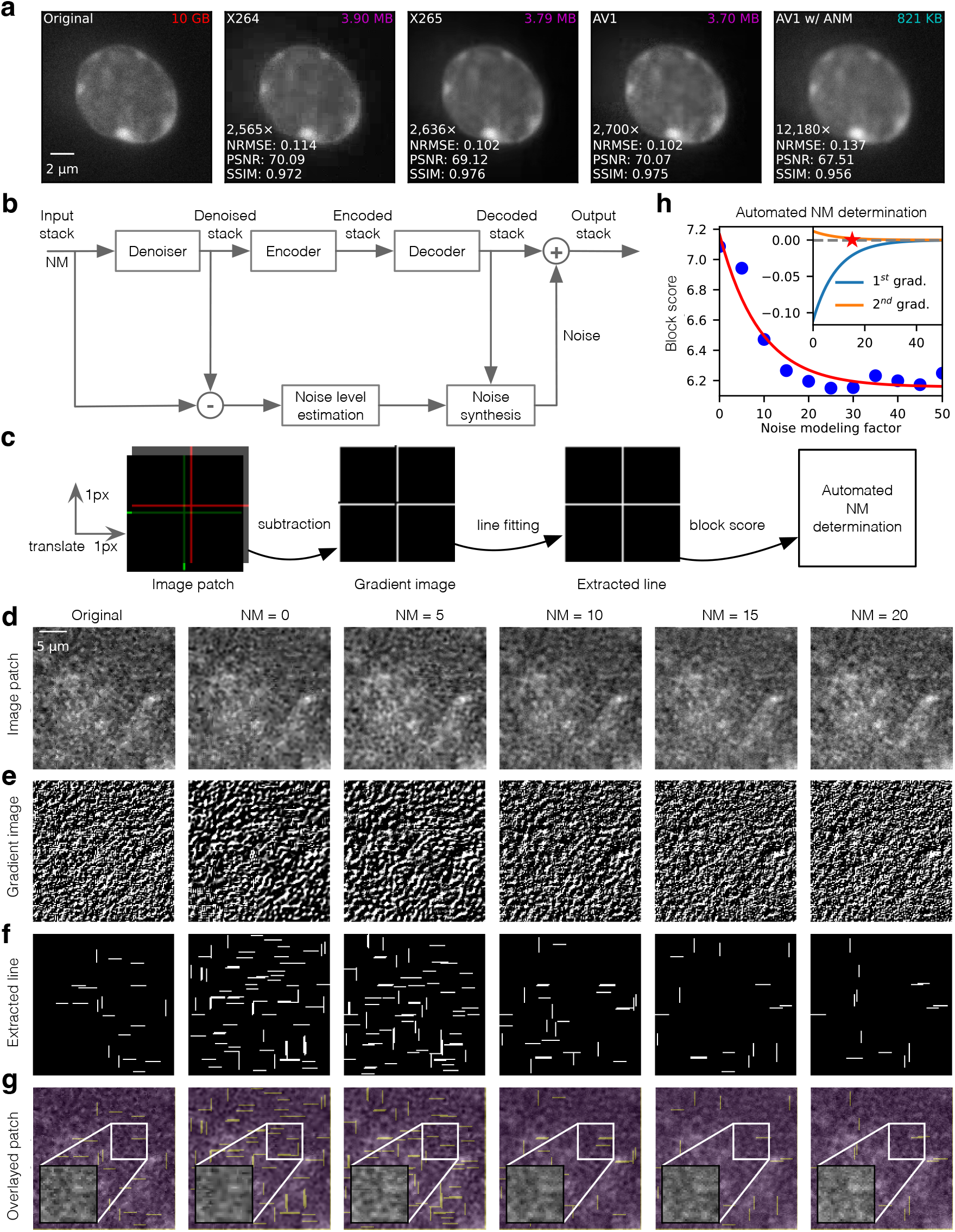
Automated determination of optimal AV1 noise modeling parameter. **a**, A magnified image of DRAQ5 stained nucleus compressed using different strategies. With appropriate AV1 noise modeling, the block artifact is minimized, enabling a compression ratio of *>*10,000 fold. **b**, Schematic of SVT-AV1 noise modeling and synthesis. A single integer value noise modeling factor (NM) can be set to control how aggressively the denoising and noise synthesis are implemented. **c**, Schematic showing each step of computing the block score for automated optimal NM determination. The final block score is defined as the log value of the total pixel number of all extracted lines. **d**–**f**, Example images corresponding to each block score computation step (image patch, gradient image, and extracted line). A noisy background image patch is shown here. **g**, Image patch overlayed with extracted lines. Magnified images in the white box visually confirmed the effectiveness of removing the block artifact by SVT-AV1 noise modeling. **h**, Automated NM determination. We fit an exponential function using block scores computed from images compressed with different NM values. The optimal NM = 15 is chosen as the tuning point of the second-order gradient of the fitted function (**h**, inset), which eliminates the block artifact while not overly smoothing the image.

Aiming to automate the NM estimation, we observed that larger NM values lead to less obvious block artifacts while producing blurrier compressed images. Thus, we hypothesize that the optimal noise modeling should use the smallest NM value that creates the minimal block artifact. Fig. 4c shows our computational pipeline to estimate a block score for determining the optimal NM value. In brief, we randomly crop out multiple sub-volume patches from the original image stack and compress them with NM across the whole allowable range (Fig. 4d). To emphasize the horizontal and vertical lines that form the block artifact in each compressed patch, we calculate the gradient of *x* and *y* dimensions by translating the image patch by 1 pixel in both dimensions and subtract this translated patch from the original patch (Fig. 4e). Next, we apply a probabilistic Hough line transform^48^ to the gradient image to extract vertical and horizontal edge lines (Fig. 4f). Finally, we sum up the number of pixels that comprise the extracted lines, as it reflects how blocky the image is and apply a log transform to the sum as the corresponding block score.

The optimal NM should result in a compressed image with the most similar block score to the original image (Fig. 4g). We iterate each NM value and plot the corresponding mean block score of all sub-volume patches (Fig. 4h). Then we fit an exponential function *ae*^−*bx*^ + *c* to the plot and determine the optimal NM as the turning point of the second-order gradient (Fig. 4h, inset). This approach determines the smallest NM that is strong enough to eliminate the block artifact while preserving sharper details.

### High-fidelity segmentation of highly compressed 3D fluorescence microscopy images

3D image segmentation is one of the most important image processing in quantifying light microscopy data^49^. This voxel-level foreground vs. background classification is extremely sensitive to changes in voxel intensity transitions. Inspired by beta10-AV1’s ability to compress images with very high fidelity of the voxel value, we examined the impact of its image size reductions on segmentation predictions using a broadly adapted machine learning model, 3D nnUNet^50^. To do so, we prepared a mouse brain section following the miriEx protocol^51^, stained the nuclei with DRAQ5, and acquired high-resolution, high signal-to-noise ratio (SNR) 16-bit line confocal microscopy images with a relatively uniform background (Fig. 5a). We then created human-annotated segmentation training datasets using a subset of raw images (Fig. 5a) and trained a 3D nnUNet model with these raw images. This pre-trained model was then used to predict the rest of the raw images as nuclei segmentation ground truth (Fig. 5b). We then overlay segmentation ground truth to segmentation from 5,880-fold, beta10-AV1 compressed images (Fig. 5c), and to segmentation from 12,180-fold, beta10-AV1 with automated noise modeling (beta10-AV1-ANM) compressed images (Fig. 5d). Compared to beta10-AV1, beta10-AV1-ANM achieved a higher segmentation agreement with the ground truth while compressing to a much higher ratio (Fig. 5e-f). For instance, 94.5% of the nuclei segmentation from beta10-AV1 compressed images has a centroid coordinate that is within a distance of *<*1 µm to the centroid coordinate of the corresponding nuclei segmentation ground truth, while this truth positive rate (TP) is 97.2% for nuclei segmentation from beta10-AV1-ANM compressed images (Fig. 5e-f, TP). Next, we plot the distance between the centroids of the corresponding nuclei segmentation pairs and found that compression with automated noise modeling significantly improves the segmentation fidelity as at least 86% of the centroids are located within a 0.2 µm distance from the ground truth, while that falls to 81% without noise modeling (Fig. 5g-h).

**Fig. 5.**
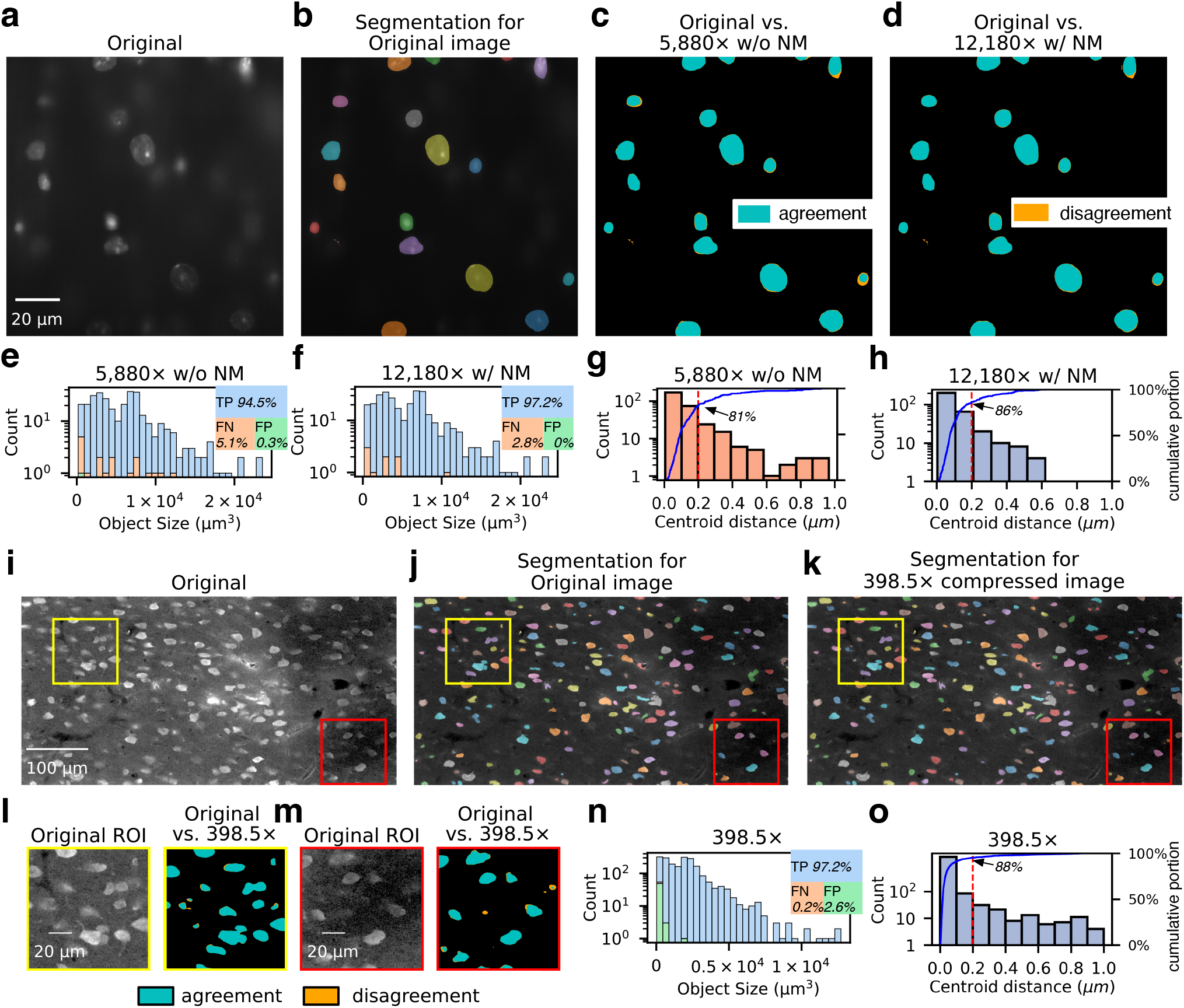
High-fidelity segmentation of highly compressed 3D fluorescence microscopy images. **a**, Single confocal image taken from a 3× expanded, DRAQ5 stained mouse brain section. **b**, Human annotation-trained 3D nnUNet was used to predict the uncompressed, original confocal images as the segmentation ground truth. Segmentation ground truth overlaid to segmentation from beta10-AV1 compressed images (**c**) and to segmentation from beta10-AV1-ANM (automated noise modeling) compressed images (**d**). **e** and **f**, Size distributions of DRAQ5 nuclei segmented from beta10-AV1 and beta10-AV1-ANM compressed images that are categorized as true positive (TP), false positive (FP), and false negative (FN), respectively. **g** and **h**, Histogram and cumulative proportion plots of centroid distances between ground truth DRAQ5 nuclei segmentations and their corresponding segmentations in the beta10-AV1 and beta10-AV1-ANM compressed images, respectively. **i**, Single confocal image taken from a NeuN stained rat brain section^28^. **j** and **k**, Human annotation-trained 3D nnUNet was used to predict neuron soma segmentations from the original confocal images and the beta10-AV1-ANM compressed images, respectively. **l** and **m**, Magnified yellow and red box regions in (**i**), respectively. Left panels, original confocal images. Right panels, overlays of segmentations from original images and from beta10-AV1-AMN compressed images. **n**, Size distributions of NeuN soma segmentations in beta10-AV1-ANM compressed images that are categorized as TP, FP, and FN. **o**, Histogram and cumulative proportion plots of centroid distances between NeuN soma segmentations in the original images and their corresponding segmentations in beta10-AV1-ANM compressed images.

Similarly, we examined the performance of beta10-AV1-ANM in compressing images with a relatively low SNR and a non-uniform background. We used a previously published dataset^28^, in which confocal images of NeuN-stained rat brain sections (Fig. 5i) were segmented manually as training soma segmentation (data not shown). We then trained a 3D nnUNet model and used it to predict NeuN soma segmentations from the rest of the original images (Fig. 5j) and from 399-fold, beta10-AV1-ANM compressed images (Fig. 5k). Close inspection of the segmentation overlays in two regions of interest (ROI) indicates a very high degree of agreement, while the false positive and false negative segmentations are mostly very small (Fig. 5l-m). Therefore, we set a threshold of 3 µm and filtered out segmentation objects that are smaller than this threshold. Again, quantification of the centroid distance between the corresponding soma segmentation pairs found in the original images and the compressed images showed that 97.2% of them are truth positive (Fig. 5n, TP) and 88% of the centroid pairs fall within a 0.2 µm distance (Fig. 5o). Finally, we computed the Dice scores for different levels of compression, which measure the agreements between two corresponding objects segmented in raw or compressed images (Supplementary Fig. 6). For NeuN and DRAQ5 images, beta10-AV1-ANM can achieve ∼95% agreement with *>*400× and *>*10,000× compressed images, respectively, indicating that it is a superior compression strategy for high-fidelity machine learning segmentation.

### High-fidelity segmentation of highly compressed electron microscopy images

EM images have extremely high content and a high background, making them notoriously difficult to compress^9^. In addition, as EM images are normally taken in very large dimensions to accommodate the very high spatial resolutions, they are normally saved as 8-bit images to reduce the file size. Therefore, we set to challenge the performance of beta8-AV1-ANM by using it to compress a small subset of the 1 mm^3^ human cortex H01 sample^52^, which has even higher noise than other EM images due to its fast 200 nanosecond scanning (Supplementary Fig. 7a-c). Close inspection of a magnified ROI (Supplementary Fig. 7d, left column), we found that AVIF (Q = 51), the 2D AV1 implementation without film grain synthesis, generated severe block artifacts at ∼ 8× compression (Supplementary Fig. 7d, middle column), in agreement with what was found in Minnen et al.^9^. To improve the compression ratio, Minnen et al. denoised the fast-scanning images by training a machine learning model with low-noise, slow 2000 nanosecond scanning images. However, this strategy requires specific training to handle images taken from different experiments. On the contrary, SVT-AV1’s film grain synthesis function utilizes Wiener type filter^53^ denoising and multi-Gaussian noise modeling that can be universally applied to all imaging conditions and produce predictable results. Indeed, we found that beta8-AV1-ANM (Q = 45, NM = 91) compression yielded block artifact-free and detail-preserved results with 14.4× compression (Supplementary Fig. 7c, right column). To be noted is that, as EM produces negative contrast, we inverted the image intensity to match fluorescence-like contrast before quantization and compression.

To evaluate how beta8-AV1-ANM compression affects deep learning model performance, we downloaded a Drosophila brain EM dataset and its human-annotated boundary map (Fig. 6a) used in the CREMI segmentation challenge^54^ and utilized them to train multiple UNet model variants^55–57^. The trained models were then used to predict boundary maps and create segmentations from the original and compressed images (Fig. 6a). Visual inspection indicates that, compared to the human annotation (Fig. 6b), 2D UNet generates almost identical boundary map and segmentation results from the uncompressed image or compressed images, including making the same mistake mistake (Fig. 6c-f, arrowhead). We further used several UNet variants to quantify the errors introduced by lossy compression. To do so, we used segmentation results from the original images as the baseline and quantified the ARAND error (whether a pixel belongs to the same cluster), the under-segmentation error (e.g., when two clusters merge into one cluster), and the over-segmentation error (e.g., when one cluster splits into two clusters). All metrics reported in Fig. 6g-i, are in the form of 1-Error, reflecting the agreement to segmentations of the uncompressed image from different compression ratios. We found that while all UNet variants create highly faithful segmentation results even using 107× compressed images, 2D UNet is the least sensitive to lossy compression. Taken together, beta8-AV1-ANM can be used for high-fidelity segmentation from highly compressed EM images.

**Fig. 6.**
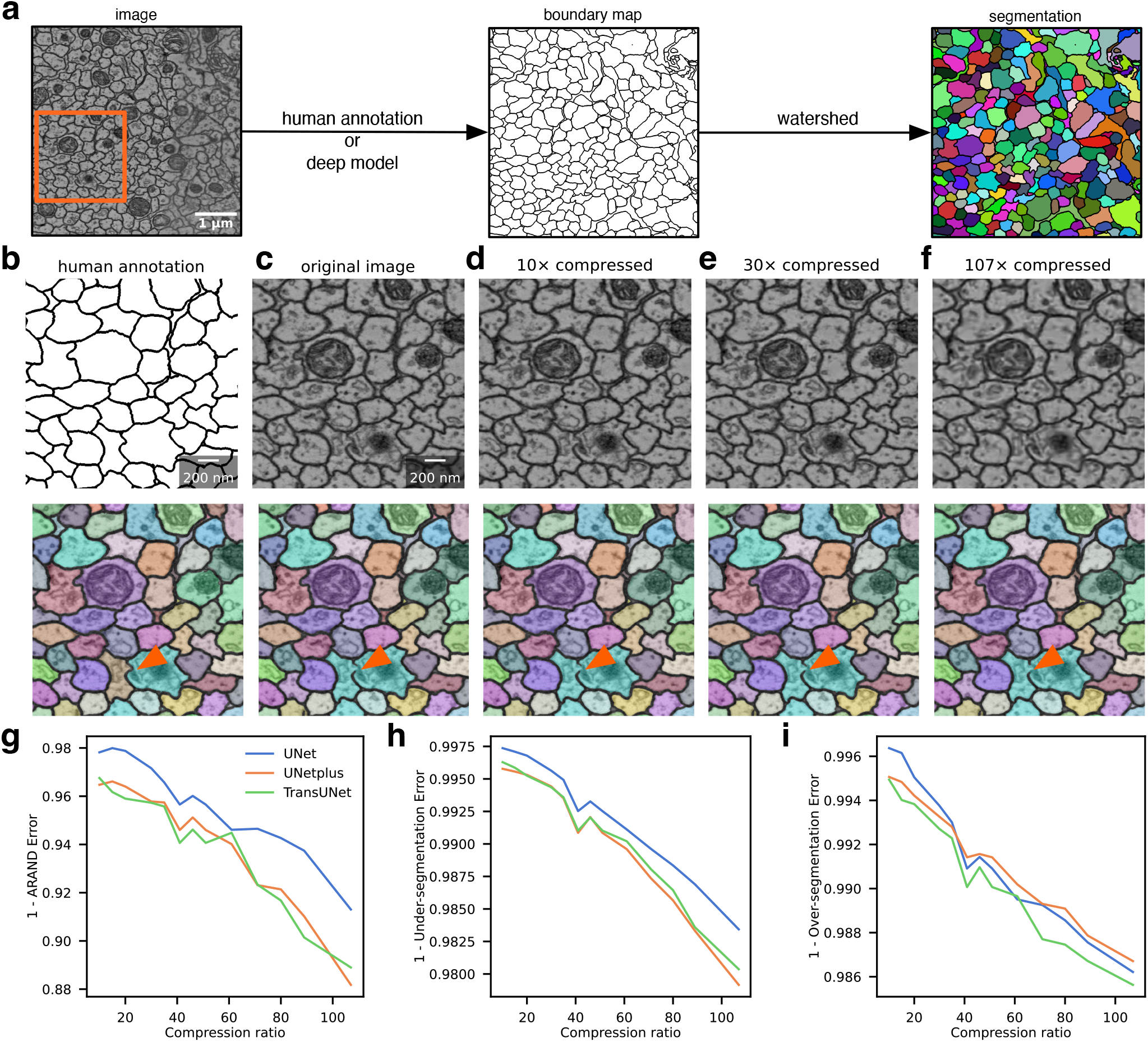
High-fidelity segmentation of highly compressed electron microscopy images. **a**, In the CREMI segmentation challenge^54^, a drosophila brain EM image dataset (left) was paired to the human-annotated boundary map (middle) to train machine learning models for cell boundary predictions. The predicted boundaries are then watershed processed to generate segmentations (right). **b**-f, Magnified views of corresponding boundary map or EM images (top row), and EM image overlaid segmentation results (bottom row) of the orange box region in (**a**). From left to right, columns are human annotation (**b**), results obtained from the original images (**c**), and 10× (**d**), 30× (**e**), or 107× (**f**) compressed images. Using three UNet variants, segmentations from the 107× compressed images were compared to the segmentations from the original images to quantify errors introduced by lossy compression. **e**-**g**, ARAND error (whether a pixel belongs to the same cluster), under-segmentation error, and over-segmentation error, presented in the form of 1 −*Error*, respectively.

## Discussion

We systematically studied image quantization and state-of-the-art video compression algorithms to understand the trade-off between the loss of information and data size reduction for light and electron microscopy image analysis. To avoid photobleaching and phototoxicity, biomedical images are commonly captured with low intensity and high noise that does not occupy the whole dynamic range allowed by modern 16-bit detectors. We developed a beta10-AV1-ANM lossy compression strategy optimized for such imaging conditions, which includes an optimal quantization step to maximally preserve entropy, followed by SVT-AV1 compression with automated parameter optimization to minimize compression artifacts.

In the quantization step, preserving as much entropy in the low-intensity regions as possible is desired. We achieve this goal by adapting a generalized power function that can effectively redistribute information coding capacity toward the low-intensity regions while reducing the total bit-depth. We also take advantage of the film grain synthesis feature of the SVT-AV1 codec to achieve a high compression ratio while eliminating the block artifact. To obtain an objectively optimal compression result, we developed an algorithm to automatically determine the noise modeling parameter. Combining these two, our beta10-AV1-ANM lossy compression pipeline empowers very high-ratio image size reduction while preserving faithful visual presentation at the voxel level and maintaining quantitative analysis fidelity.

Our results show that high-fidelity beta power quantization results in better analysis accuracy after lossy compression in noise-sensitive applications, such as fluorescent spot localization for SMLM. This is an analogy to a dual-stage amplification experiment, such as single-cell mRNA sequencing library preparation, in which choosing a high-fidelity linear amplification instead of an error-prone exponential PCR as the first stage amplification can generate a higher quality library for gene recovery^58^. Automated optimization for SVT-AV1 film grain synthesis further enables the image to be compressed to orders of magnitude higher compression ratio for object segmentation in light and electron microscopy images. Our beta10-AV1-ANM’s ability to compress volumetric images to ten thousand folds while preserving accurate segmentation results is exciting and sheds light on managing the ever-growing image data size generated by modern microscopy.

That said, better and faster noise reduction and modeling algorithms are highly desired to replace the default SVT-AV1 implementation. In parallel, deconvolution algorithms can also be applied prior to noise reduction to increase the compression ratio by reducing image entropy, i.e., the information density of the image. To add realistic noise back to the denoised images, SVT-AV1’s multimodal Gaussian noise modeling is versatile and effective for all the images we have tested, including those taken by EM. It is worth noting that our study focused on evaluating the impact of image compression when using analysis methods and machine learning models developed with uncompressed images. We expect that if these models were retrained using the compressed images, the performance would be even more impressive. In addition, as our compressed images are visually very similar to the uncompressed ones, we expect minimal impacts on human annotations, including generating segmentation and neuron tracing ground truth for machine learning training purposes.

Finally, to encourage community adaptation of the most up-to-date compression codecs, we developed forward-compatible APIs with the FFMEPG^59^ library. We implemented the beta10-AV1-ANM algorithms in a C-based HDF5^60^ filter, which can be used in Python programming via hdf5plugin^61^ and h5py^62^. We also developed a Java-based ImageJ/Fiji^19^ plugin to allow compression, decompression, visualization, and manipulation of terabyte-scale images using the virtual stack interface. These plugins also support hardware-accelerated video codecs, such as those implemented in the NVIDIA CUDA framework. Together, these tools will form a basis for working with high-ratio, lossy compressed images in existing pipelines.

## Online Methods

### Data collection

Images were collected from their respective original publications (See **Data availability**). For the DRAQ5 images, C57/B6 mouse brains were sliced coronally into 100 µm sections and expanded to ∼ 3× to the miriEx protocol^51^, followed by staining with 5 mM DRAQ5 solution (ThermoFisher Scientific). Z stacks images were collected with a 500 nm step size using a custom-built line confocal microscope excited by a 642 nm laser and preprocessed using custom Python scripts^63^. All animal studies were carried out under the supervision of a protocol approved by the University of Michigan Institutional Animal Care & Use Committee (IACUC).

### Quantization

#### Linear quantization

We first divide the raw image intensity by its brightest intensity and then multiply it by the highest value that the low-bit image can represent, i.e., 2^8^− 1 = 255 for an 8-bit image and 2^10^− 1 = 1023 for a 10-bit image.

#### Square root quantization

The intensity of each voxel in a 16-bit image was taken by square root to convert it into an 8-bit value.

#### Beta quantization

A generalized power function

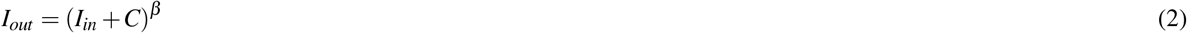

was used to determine an optimal *β*, for beta quantization, where *I*_*out*_/*I*_*in*_ is intensity after/before quantization, and *C* is a constant, which in our case equals 0. We first identify the maximum intensity *I*_max_ in the original image and then use

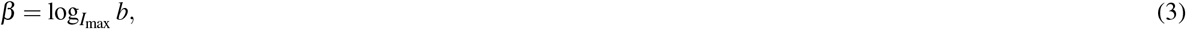

where *b* = 255 if transforming to an 8-bit image, and *b* = 1023 if transforming to a 10-bit image. In practice, one can consider using, e.g., the 98% maximum intensity value to avoid hot pixel errors caused by shot noise or detector defects.

### Image variance and entropy

Variance was calculated as

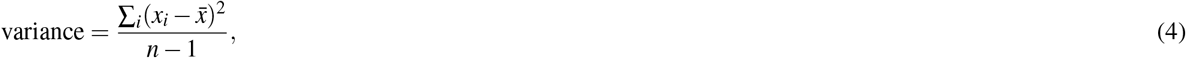

where within a sliding square window of side length 5, thus *n* = 25, *x*_*i*_ is the intensity of each pixel, 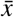 is the mean intensity of all pixels.

Relative variance error was calculated as the variance difference between two comparing images that have been normalized to the intensity of the pixel in the raw image im_raw_ as

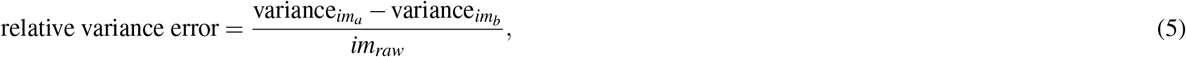

quantifying the relative variance error of *im*_*a*_ and *im*_*b*_ from the perspective of the raw image.

Entropy, as know as Shannon Entropy^25^ was calculated as

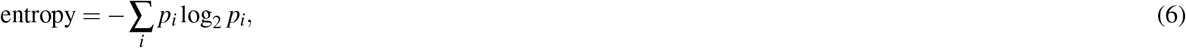

where *p*_*i*_ represents the percentage of i-th bin in the normalized histogram under the sliding square window of side length 5. The entropy difference was calculated between the 16-bit raw image and the decompressed and 16-bit reverse-transformed image with different quantization strategies.

### Compression algorithm benchmark

#### Computation environment

A server with two AMD EPYC 7262 8-Core processors and 256GB RAM was used to build a virtual `Python V3.11.3` environment under `Anaconda V23.1.0`^64^. `hdf5plugin V4.0.1`^61^ and `h5py V3.8.0`^62^ were used for testing the HDF5 plugin, which was developed in C. All files were saved into a memory buffer rather than a disk to ensure that disk latency or bandwidth did not impact benchmark results. The compression ratios were calculated based on the size of storing the uncompressed data into the memory buffer. For compression tests, default settings on thread numbers for parallelization were used since most video compression algorithms are well-tuned to support multi-thread encoding.

#### Compression algorithms

Blosc-Zstd was set to a compression level of 5 in hdf5plugin to achieve the highest size reduction. H.264^16^ and H.265^17^ were implemented using the X264^20^ (version: `0.164.x`) and X265^21^ (version: `3.5+1-f0c1022b6`) libraries, respectively. AV1^18^ was implemented using the SVT-AV1^22^ (version: `1.4.1`) library. All codecs were built using `GCC 10.3.0` for a 64-bit Linux environment (`Ubuntu 22.04.1 LTS`).

#### Compression quality comparison metrics

Root Mean Squared Error (RMSE) is defined as

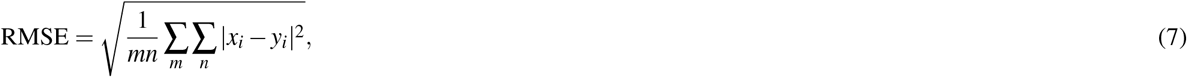

where *m, n* are numbers of pixels, *x, y* are images, and *i* is the index for a pixel. Normalized root mean squared error (NRMSE) is defined as

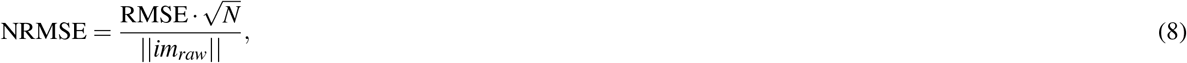

where || · || denotes the Frobenius norm and *N* = size(*im*_*raw*_). Peak signal-to-noise ratio (PSNR) is defined as

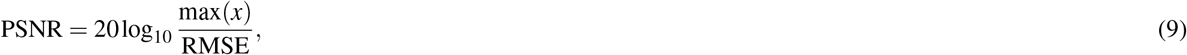

where max(*x*) represents the highest intensity of the data, e.g., 255 for 8-bit image, measuring the sharpness of the modified image. Structural similarity index measure (SSIM) predicts the perceived quality of a modified image, and is used for measuring the similarity between two images. SSIM needs to slide a small window, in our case, size of 7 pixels, throughout both images, and is calculated as

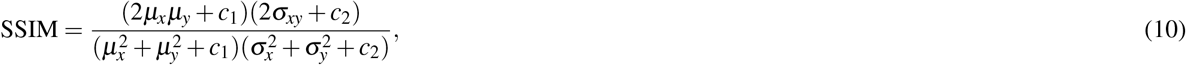

where *x* and *y* are corresponding windows from two images, *µ*_*x*_/*σ*_*x*_, *µ*_*y*_/*σ*_*y*_ is the mean/variance of *x* and *y, σ*_*xy*_ is the covariance of *x* and *y, c*_1_ and *c*_2_ are two constants to stabilize the division. The final SSIM index is obtained by averaging over values from all sliding windows. We use implementations from `scikit-image V0.20.0`^65^ for all metrics.

### Single-molecule localization

Fluorescent spot identification and fitting were obtained using `ThunderSTORM`^37^ (`dev-2017-01-28-b1`) in Fiji. True Positive (TP), False Positive (FP), and False Negative (FN) were obtained by comparing localization results between the original images (ground-truth, GT) and compressed images (prediction). TP is defined as the points in GT found in the prediction within a distance threshold of 320 nm. FP denotes points that are not in GT but found in the prediction, while FN represents points that are in GT but not found in the prediction. The distance threshold is defined as the largest Euclidean distance to consider whether two located points are a pair or not, which is set to 2 times of pixel resolution (160 nm). Thus,

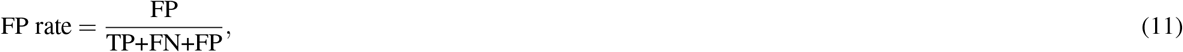

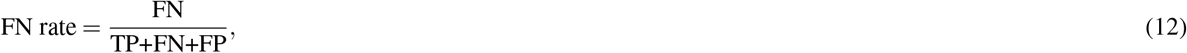

reported in percentage. The spot identification error rate (SIER) is defined as

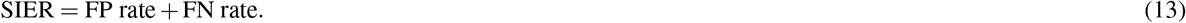

The centroid localization error (CLE) is defined as the absolute Euclidean distance in nanometers between the closest pair of predicted and GT points in the corresponding dimension.

### Automated noise modeling for minimizing compression block artifacts

To acquire the optimal AV1 noise modeling (NM) factor for a pre-determined quality factor Q, we first randomly sample ten sub-volumes of size (128, 128, 2) from the original image stack and then compute their block scores for all possible noise modeling factors. The block score for each compression Q factor and NM factor is calculated by summing the total pixels in the extracted intensity gradient lines and applying a log-scale transform. Block scores for all NM factors are used to fit an exponential function

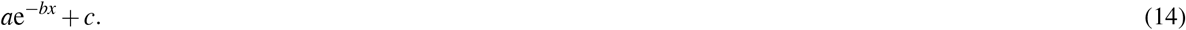

The turning point of the second-order gradient of the fitted function is chosen as the optimal noise modeling factor (Fig. 4f), which balances the strength of noise modeling and the preservation of image details. If it fails to fit a function sometimes due to the fitting algorithm, e.g., reaching maximum iterations, we can directly use gradients from the Gaussian-smoothed block score array and find the turning point of the array’s second-order gradient as an alternative.

### 3D fluorescence microscopy image segmentation

Manually annotated 108 image patches of dimension (128, 160, 320) were used as segmentation training datasets. The 108 pairs of raw images and mask images were split into 86 pairs for training and the rest 22 pairs for validation. nnUNet^50^ was trained in `Pytorch V1.10`^66^. Standard data augmentations such as rotation, scaling, Gaussian noise, Gaussian blur, brightness, and contrast adjustment, simulation of low resolution, gamma augmentation, and mirroring were used to permute the input data. Initial learning rate (lr) was set to 1*e*^−2^ and employed a *poly* decay strategy defined as

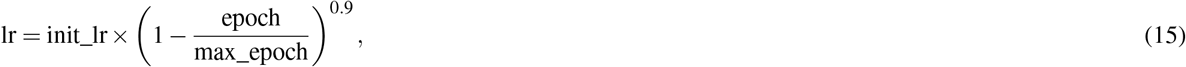

where epoch is the current epoch, init_lr is the initial learning rate, and max_epoch is the maximum training epochs. max_epoch was set to 200, and each epoch includes 250 iterations. Each epoch took 280 seconds, while the whole training process took 15.5 hours. Stochastic gradient decent^67^ was adopted as the optimizer. The momentum was set to 0.99 and the weighted decay was set to 3*e*^−5^. A combination of cross-entropy loss and dice loss was exploited to train the network as suggested in nnUNet, where the weights for two losses were equal. We utilized instance normalization as our normalization layer. The final dice on the validation dataset was 96% average during the training process. The inference time of an image with the shape (1000, 448, 448) took around 2 minutes. All experiments were conducted on a single NVIDIA RTX A5000 GPU with 24GB memory.

The network was trained with deep supervision. The combined loss (cross-entropy+dice) was used multi-scale, added in the decoder to the last three stages (the three highest resolutions). For each deep supervision output, we downsampled the ground truth segmentation mask for the loss computation with each deep supervision output. The final training objective was the sum of all resolutions losses as

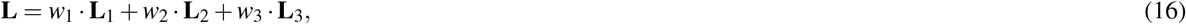

where the weights were halved by each downsampling in resolution (*w*_2_ = 0.5*w*_1_; *w*_3_ = 0.25*w*_1_, etc), and all weights were normalized to sum to 1.

For analyzing the results, we first use `scikit-image` scientific package to locate the centroid for each 3D cell given the segmentation predicted by the nnUNet. The centroid distance was the absolute Euclidean distance in pixels among different dimensions. The Dice score is defined as

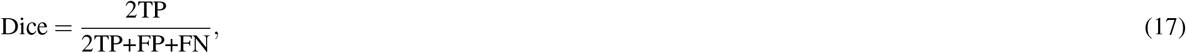

where TP is the agreement of the foreground classification between two segmentations, and FP+FN are the sum of disagreements between two segmentations. Dice measures the segmentation accuracy at a pixel level, scaling from 0 to 1 where 1 indicates perfectly agreed to the ground-truth segmentation.

### 2D electron microscopy image segmentation

The whole 3D raw images and mask images were split into the first 75 frames of pairs for training and the rest 50 frames of pairs for testing. Standard data augmentations such as Gaussian blur, motion blur, rotation, Gaussian noise, and grid distortion were utilized. The training image size was (512, 512) with a batch size of 8. The initial learning rate (lr) was set to 1*e*^−2^ and adopted a *poly* decay strategy as the same in Eq. (15), where the maximum epoch count was set to 300. Each epoch took 10 seconds, while the whole training process took 50 minutes. SGD was used as the optimizer and the momentum was set to 0.99. The weighted decay was set to 1*e*^−4^. Both cross-entropy loss and dice loss were used to train the network, where their weights were the same. The inference time of an image with the image shape (50, 1250, 1250) elapsed around 24 seconds. All experiments were conducted on a single NVIDIA RTX A5000 GPU with 24GB memory using `Pytorch V1.10`.

To analyze the fidelity of the segmentation results of compressed to that of uncompressed, we trained different UNet variants and quantitatively evaluated them using ARAND error under-segmentation error, and over-segmentation error.

ARAND error (Adapted Rand error^68^), is defined as

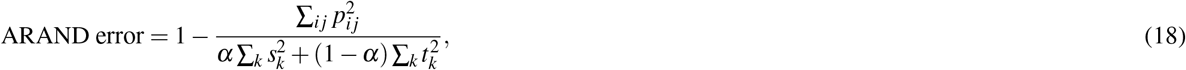

where *p*_*i j*_ is the probability that a pixel has the same label in the compressed image and in the uncompressed image, *t*_*k*_ is the probability that a pixel has label *k* in the uncompressed image, and *s*_*k*_ is the probability that a pixel has label *k* in the compressed image. By default, it weighs precision and recalls equally in the calculation. When *α* = 0, ARAND equals recall. When *α* = 1, ARAND equals precision. The under-segmentation error and over-segmentation error are defined as conditional entropy in^69^ as

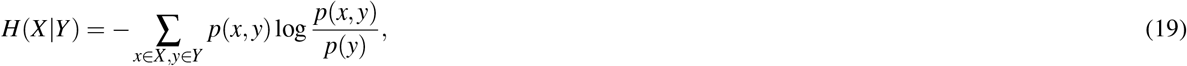

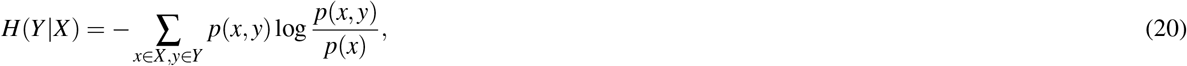

respectively, where *X* is the segmentation results of the uncompressed image, *Y* is the one of the compressed image, and *x, y* are data points, i.e. pixels. We then normalize both errors by its upper bound^69^. For all these metrics, we use the implementations from `scikit-image`.

### Video compression codec HDF5 plugin

We implemented video compression codecs in C programming language as an HDF5 plugin `HDF5 V1.14.0`^60^ and `FFMPEG V5.1.2`^59^ libraries. Documentation for installation, plugin architecture, API, and coding examples is detailed in **Code availability**). Benefiting from the underlying `HDF5` and `FFMPEG` libraries, our plugin supports auto-chunking (different chunk sizes for compression) and up to 8K resolution compression, i.e., 7,680 pixels in width and 4,320 pixels in height. Future compression algorithm updates can be conveniently accomplished by updating the FFMPEG library with a few code modifications.

### Video compression codec ImageJ/Fiji plugin

We developed an ImageJ/Fiji plugin to allow lossy compression and decompression of image stacks that can be imported into ImageJ/Fiji. To open and manipulate a compressed image, the virtual stack format is recommended due to the size of the decompressed images can often exceed the memory capacity of the user’s computer.

## Data availability

Brainbow images can be downloaded at https://www.cai-lab.org^29,70^. STORM images are from^30^, NeuN staining images are from^28^. EM images in Fig. 2 are from^31^, EM images used in Fig. 6 segmentation analysis are from^54^. Human cortex H01 EM images^52^ (Supplementary Fig. 7). Other data used in this work can be obtained through the authors upon request.

## Code availability

HDF5 plugin, ImageJ/Fiji plugin, and documentation will be made publicly available at:

- https://github.com/Cai-Lab-at-University-of-Michigan/ffmpeg_HDF5_filter

Analysis codes used to produce the results in this report are available at:

- https://github.com/Cai-Lab-at-University-of-Michigan/lossy_image_storage_2024

## Acknowledgements

This work was funded by the United States National Institutes of Health (NIH) grants RF1MH123402, RF1MH124611, and RF1MH133764 to YY and DC. LAW was supported by a University of Michigan Rackham Predoctoral Fellowship. We thank the University of Michigan Advanced Research Computing (ARC) for their assistance with high performance computing. GPU Hardware used in part of this analysis was provided by an NVIDIA academic computing grant to LAW and DC.

## Author contributions

BD, LAW, and DC conceptualized this work. BD wrote the plugin and conducted the analysis of video compression codecs. BX trained models for 3D segmentation analysis. BD and DC conceptualized the ImageJ GUI, and WJL and AL implemented the GUI, wrote documentation, and tested its functionality. LAW collected the unpublished sample data. This project was supervised by YY and DC. BD, LAW, and DC drafted this manuscript, which was edited and approved by all authors.

