## Supplemental Figures for "Artifact-Minimized High-Ratio Image Compression with Preserved Analysis Fidelity"

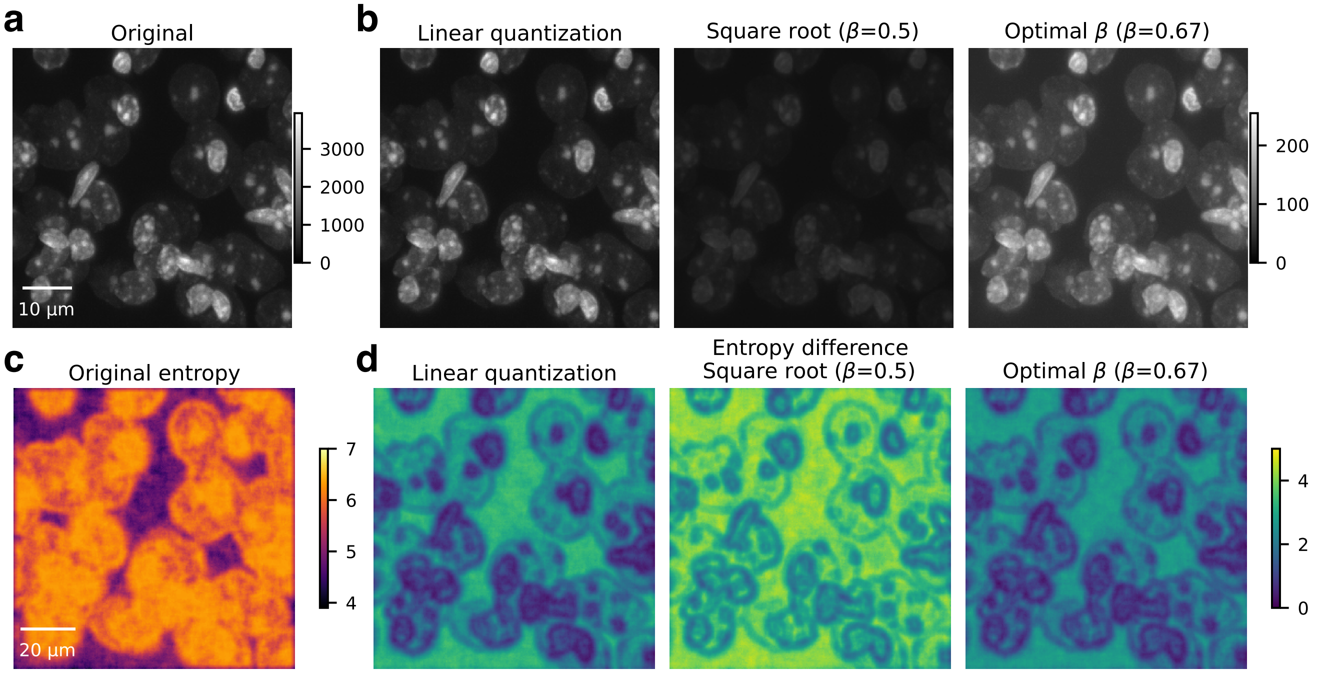


**Supplementary Fig. 1 | Optimal beta quantization has the least entropy loss compared to other quantization methods.** **a**, Original 16-bit confocal images of cell nuclei fluorescently labeled with DRAQ5 from a mouse brain section. **b**, From left to right: 8-bit image transformed by linear quantization, square root quantization ($\beta = 0.5$), and optimal beta quantization ($\beta = 0.67$) calculated based on the maximum intensity of the original image (**a**), respectively. **c**, Entropy of the original 16-bit image (**a**). **d**, From left to right: entropy differences between the original 16-bit images and the 16-bit images recovered from (**b**), respectively.

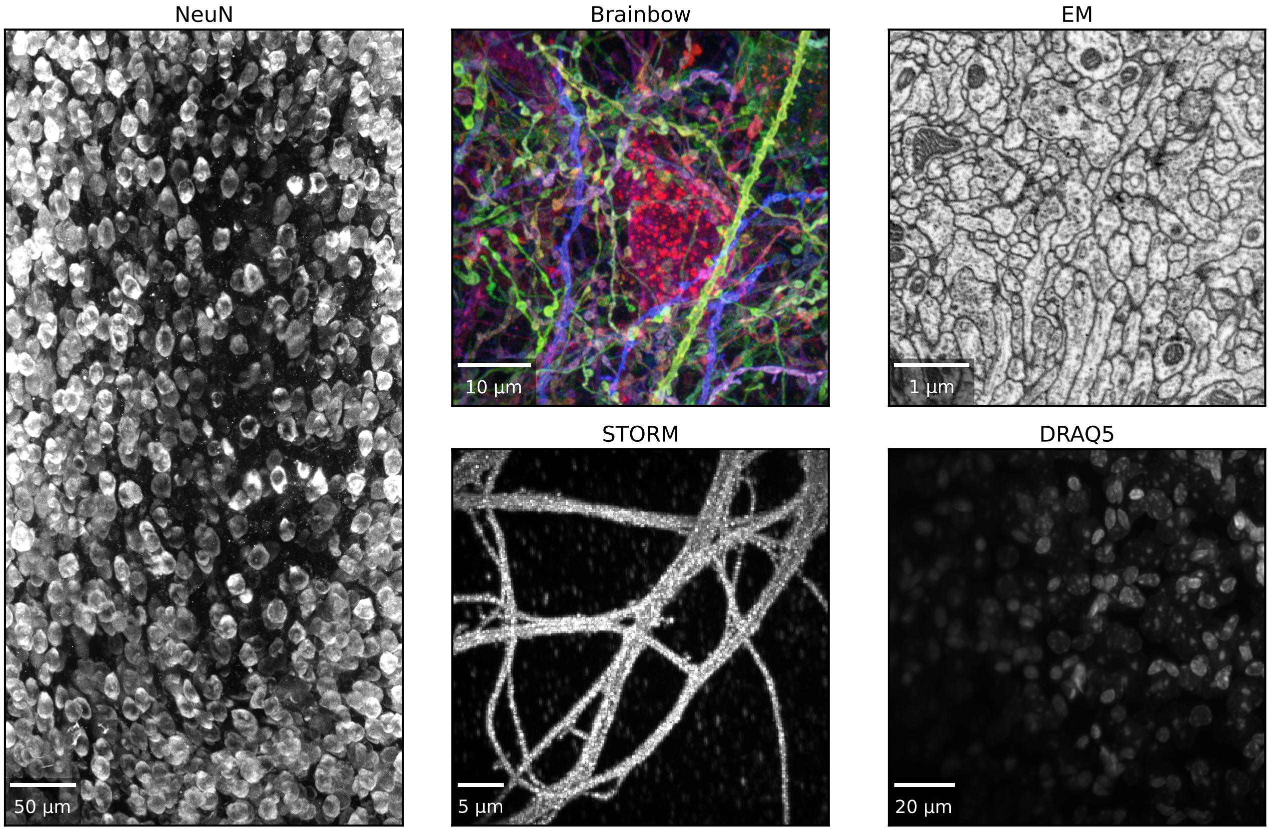


**Supplementary Fig. 2 | Overview of lossy compression benchmark image datasets.** Z-axis maximum intensity projections: NeuN, low contrast point scanning confocal microscopy images of NeuN staining[28]; Brainbow, high contrast point scanning confocal images of Brainbow-labeled mouse neurons[29]; DRAQ5, expansion line scanning confocal microscopy images of DRAQ5 stained mouse brain nuclei; and EM, Electron microscope scanned Drosophila melanogaster larval neuropil[31]. Time-axis maximum intensity projection: STORM, STochastic Optical Reconstruction Microscopy images of $\beta$2-spectrin labeling in hippocampal neurons[30].

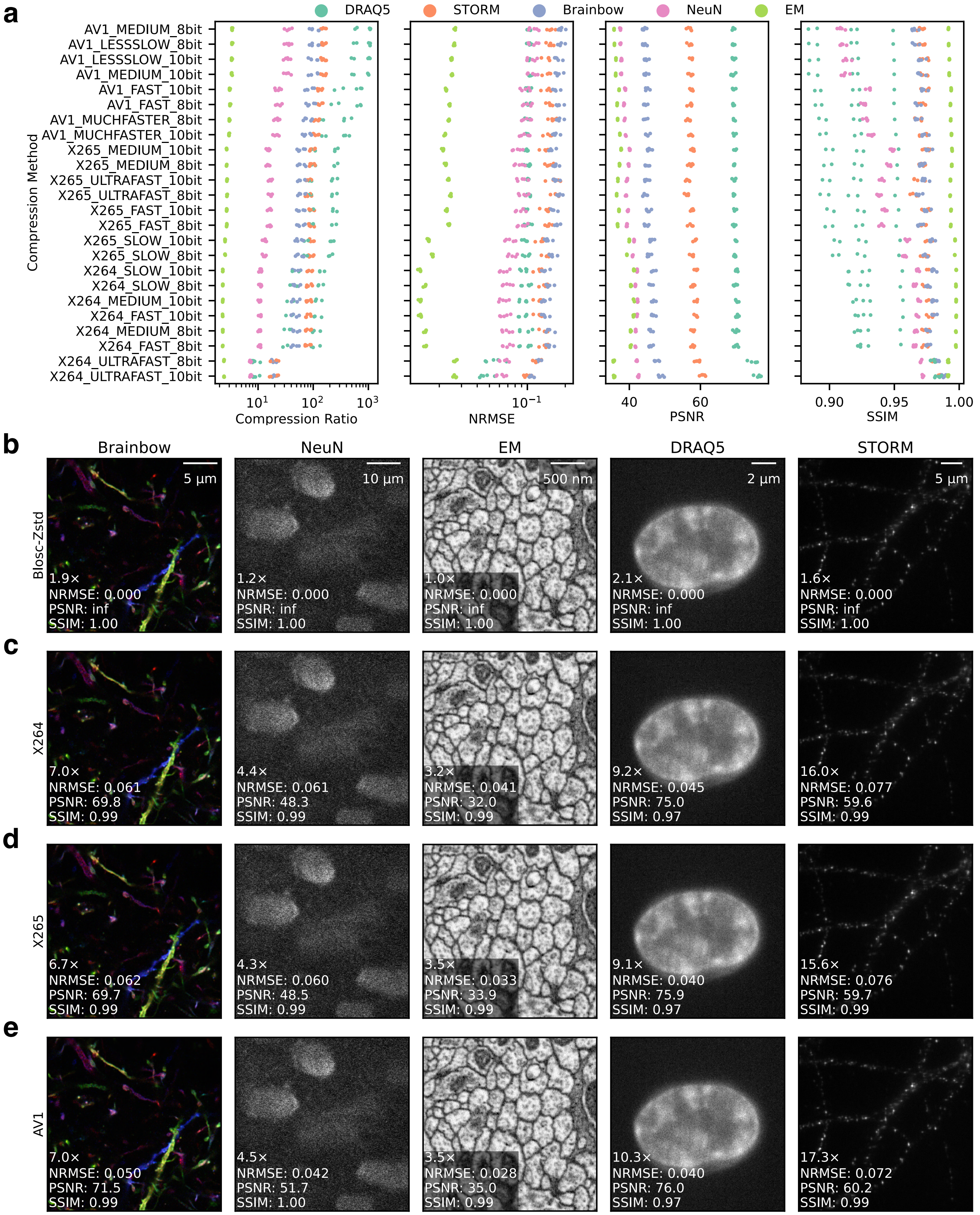


**Supplementary Fig. 3 | Comparison of Blosc-Zstd, X264, X265, and AV1 compression. a**, Quantitative comparisons between different setups for different video compression methods, sorted by descending order of compression ratio. Qualitative sample results of **b**, Lossless Blosc-Zstd compression; **c**, X264 (H264) video compression; **d**, X265 (HEVC) video compression; **e**, AV1 video compression. The compression ratio, normalized root mean squared error (NRMSE), peak signal-to-noise ratio (PSNR), and structural similarity index measure (SSIM) are reported for each compressed sample.

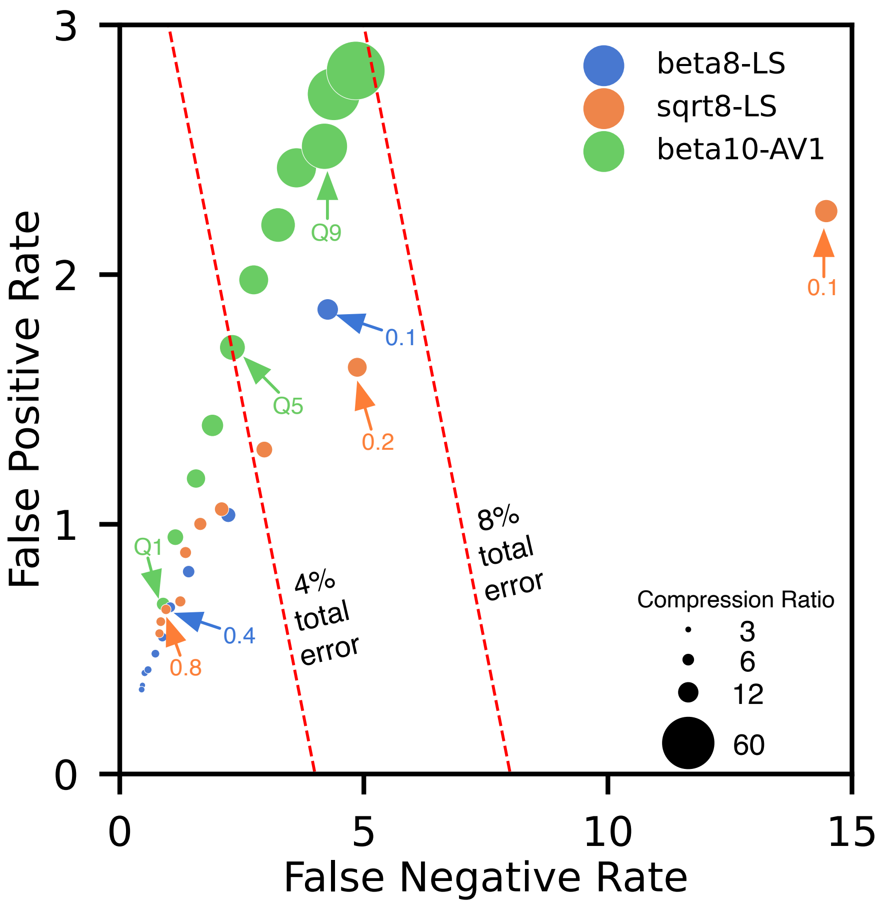


**Supplementary Fig. 4 | Plot of image compression ratio, single molecule localization false positive (FP) rate, and false negative (FN) rate using different compression setups.** The total error rate combines FP and FN rates. sqrt8-LS, linear-scaled 8-bit square root quantization followed by Blosc-Zstd lossless compression; beta8-LS, linear-scaled 8-bit optimal beta quantization followed by Blosc-Zstd lossless compression; beta10-AV1, 10-bit optimal beta quantization followed by AV1 lossy compression.

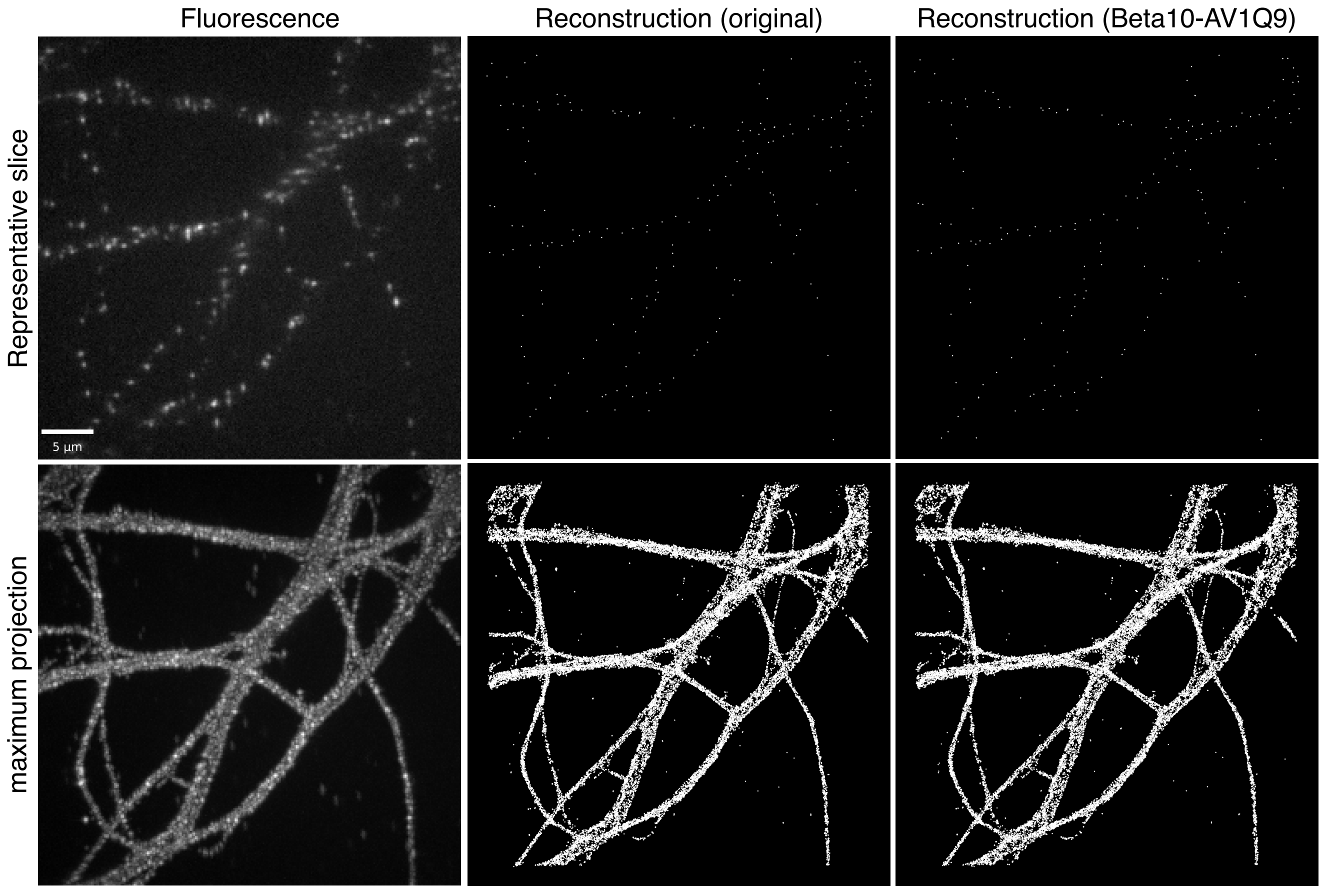


**Supplementary Fig. 5 | Super-resolution image reconstruction by single-molecule localization using original and AV1 compressed images.** Left column, original fluorescent images of $\beta$2-spectrin labeling in hippocampal neurons. Middle column, single-molecule spots localized in original images. Right column, single-molecule spots localized in 43.7× compressed images. Beta10-AV1Q9, 10-bit optimal beta quantization followed by AV1 lossy compression with quality factor 9.

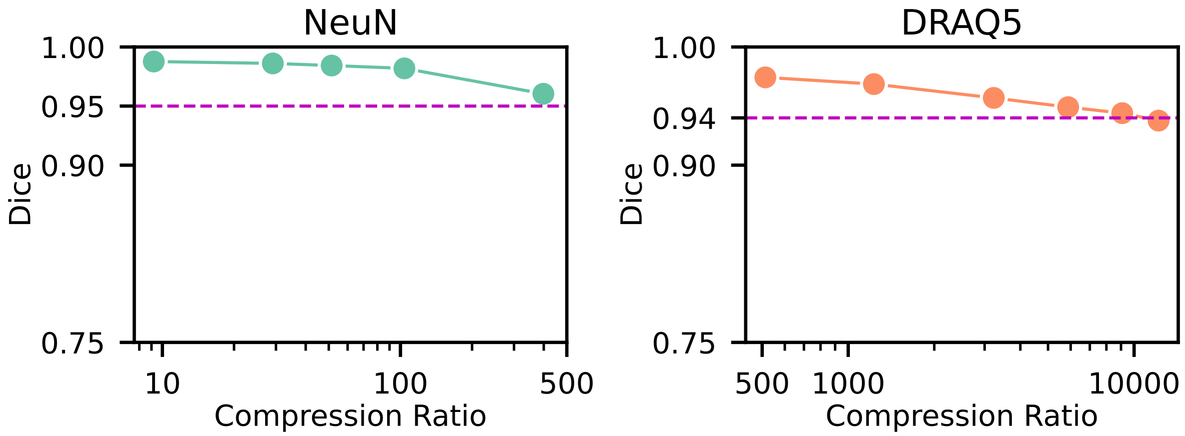


**Supplementary Fig. 6 | Dice score plots over different levels of compressions for NeuN and DRAQ5 stained rodent brain imaged by point scanning confocal and 3x expansion line scanning confocal microscopy, respectively.** Dice scores for different levels of compression measure the agreements between two corresponding objects segmented in raw or compressed images.

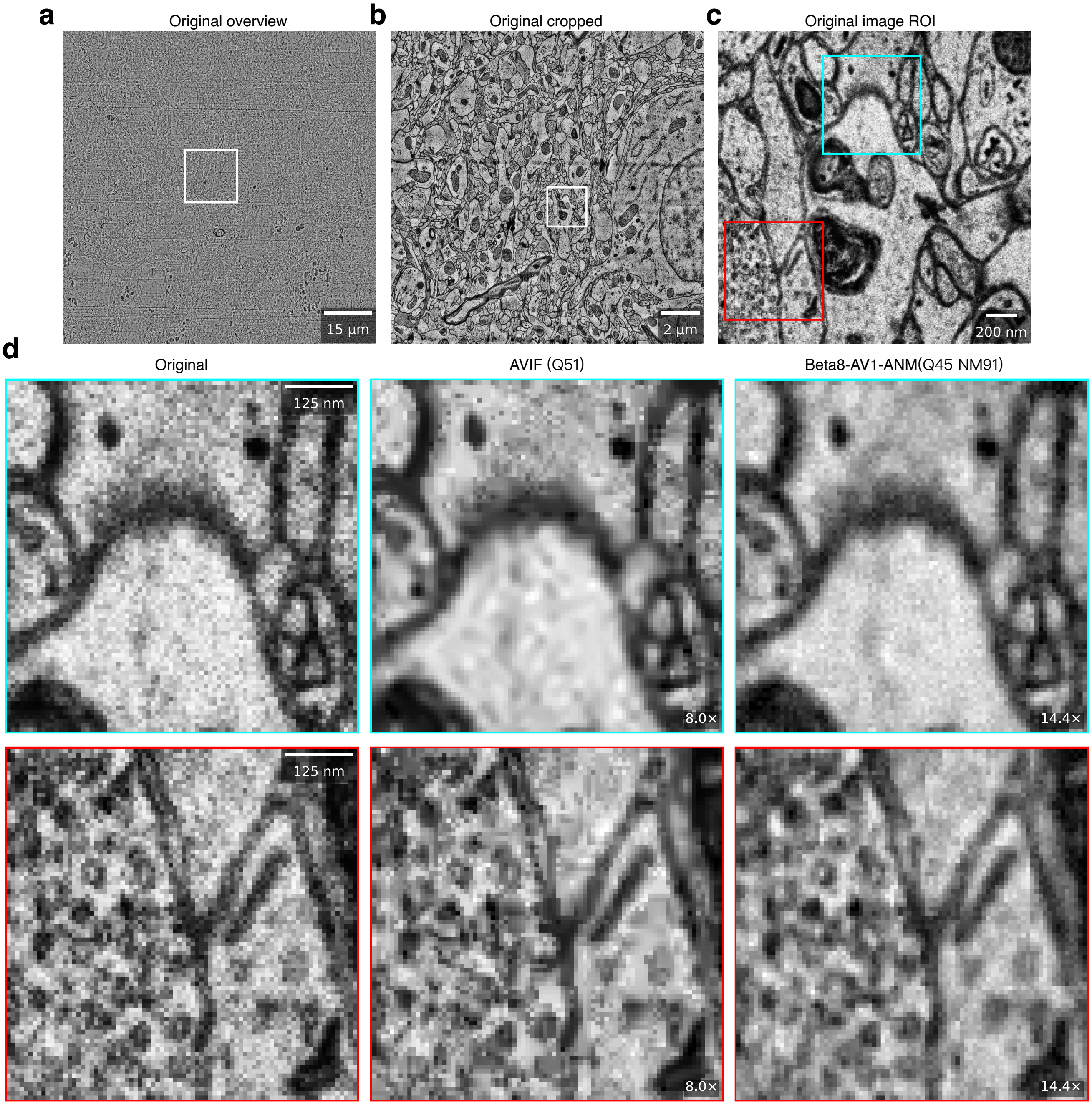


**Supplementary Fig. 7 | Beta8-AV1-ANM enables high-quality lossy compression of very noisy electron microscopy images.** **a**, Overview of the human cortex H01 dataset[52]. **b**, Magnified white box region in (**a**). **c**, Magnified white box region in (**b**). **d**, Top and bottom rows show magnified cyan and red box regions in (**c**), respectively. Left column, original images. Middle column, 2D AV1 compression (AVIF) with quality factor = 51 (Q51) generates 8.0× compressed images full of block artifacts. Right column, beta8 quantization followed by AV1 compression with noise modeling (Q45, NM91) generates 14.4× compressed images without block artifacts.
